# Link between lipid remodeling and ESCRT-III function in multivesicular body formation

**DOI:** 10.1101/2023.10.10.561655

**Authors:** Ralf Kölling

## Abstract

Despite a tremendous amount of work, it is still unclear how the endosomal sorting complex required for transport (ESCRT)-III complex acts in membrane remodeling and abscission. Here we present evidence that a change in membrane composition could be connected to ESCRT-III function during multivesicular body (MVB) formation. The central observation was a strong synergistic effect of two mutations on the turnover of an endocytic cargo protein. One mutation deletes Tms1, a yeast SERINC homologue. Human SERINC3 and SERINC5 are HIV-1 restriction factors and have been shown to act as scramblases, flipping phospholipids between membrane leaflets. The other mutation deletes the Vps68 subunit of the Vps55/Vps68 complex, which loosely resembles Tms1 in its overall structure. The strong synergistic effect suggests that Tms1 and Vps55/Vps68 perform a similar function. Since we could also show that Vps55 and Tms1 physically interact with ESCRT-III, we propose that a scramblase is recruited to ESCRT-III and plays a role in intraluminal vesicles formation at MVBs.

## Introduction

The endosomal sorting complex required for transport (ESCRT)-III complex is involved in a plethora of membrane remodeling processes in eukaryotic cells. Despite a tremendous and impressive amount of work, it is still unclear how exactly ESCRT-III performs its function (Henne *et al*. 2013; Gatta and Carlton 2019; Remec Pavlin and Hurley 2020; Vietri *et al*. 2020; Pfitzner *et al*. 2021). There are eight basic types of ESCRT-III proteins with some isoforms in higher eukaryotic cells; four of these (Snf7, Vps2, Vps20 and Vps24 in yeast) are considered to constitute ESCRT-III (Babst *et al*. 2002). The other members of this protein family are called ESCRT-III associated or ESCRT-III like (Nickerson *et al*. 2006; Azmi *et al*. 2008; Rue *et al*. 2008). The current view is that ESCRT-III proteins form spiraling filaments on membrane surfaces that in conjunction with the AAA-ATPase Vps4 exert mechanical force onto the membranes, which lead to their deformation and to scission events.

We have been especially interested in the ESCRT-III associated protein Mos10 (also known as Vps60) (Kranz *et al*. 2001). Recent evidence suggests that Mos10 forms an alternative ESCRT-III complex independent of the canonical Snf7 containing complex (ALSLEBEN AND KÖLLING 2022; Pfitzner *et al*. 2023). Both complexes appear to be involved in the formation of intraluminal vesicles (ILVs) during multivesicular body (MVB) formation in the endocytic pathway (Pfitzner *et al*. 2023).

Recently, we identified Vps68 as an interaction partner of Mos10 (ALSLEBEN AND KÖLLING 2022). Vps68 forms a complex with Vps55, whose role in endocytic trafficking has been studied previously (Belgareh-Touze *et al*. 2002; Huh *et al*. 2003; Schluter *et al*. 2008; Suzuki *et al*. 2021). Both proteins have a similar structure basically consisting of four transmembrane spanning segments (TMDs). Deletion of *VPS55* and/or *VPS68* has only a moderate effect on the degradation of endocytic cargo proteins (Schluter *et al*. 2008; ALSLEBEN AND KÖLLING 2022). But here, we identified another factor (*TMS1*), whose deletion leads to a complete block in the degradation of an endocytic cargo protein in combination with the *VPS68* deletion. This strong synergistic effect suggests that the Vps55/Vps68 complex and Tms1 perform a similar function.

Tms1 is a yeast member of the SERINC (serine incorporator) protein family, a group of highly conserved membrane proteins with ten membrane spanning segments (Pye *et al*. 2020; Leonhardt *et al*. 2023). In humans there are five members of this protein family, SERINC1-5. Yeast Tms1 shows a remarkably high sequence identity of 33 % with human SERINC1. This high degree of conservation suggests that SERINCs perform a central function in eukaryotic cells. Initially it was thought that SERINC facilitates the incorporation of serine into phosphatidylserine and sphingolipids (Inuzuka *et al*. 2005), but in a later study no significant effect of SERINC expression on the lipid composition of producer cells could be detected, arguing against the initially proposed function (Trautz *et al*. 2017). SERINC3 and SERINC5 are HIV-1 restriction factors (Rosa *et al*. 2015; Usami *et al*. 2015). The presence of these proteins in the envelope of viral particles reduce HIV-1 infectivity by up to 100-fold. Normally, the HIV-1 Nef protein prevents the incorporation of SERINCs into viral particles by triggering their endocytosis. Recently, evidence has been presented that SERINCs are scramblases (Leonhardt *et al*. 2023), non-ATP dependent lipid transporters, which flip phospholipids between the leaflets of the lipid bilayer.

Cryo-EM structures of human SERINC3 and SERINC5 and its *Drosophila* orthologue have been presented (Pye *et al*. 2020; Leonhardt *et al*. 2023). The ten TMDs of SERINC5 are arranged into two α-helical bundles consisting of four TMDs each, which are connected by an asymmetric cross of two TMDs. Vps55 and Vps68 consist of four TMDs each. Thus, it is conceivable that Vps55/Vps68 could be a distant, fragmented member of the SERINC family. There are no obvious sequence identities between Vps55/Vps68 and Tms1, but a relatedness may be difficult to prove. This is exemplified by Ice2, which was recently identified as another SERINC member in yeast, only after extensive application of sophisticated bioinformatic tools (Alli-balogun and Levine 2021).

Our study establishes a link between lipid remodeling and ESCRT-III function. We show that the putative scramblase Tms1 and Vps55/Vps68 directly associate with ESCRT-III. An alteration of the lipid composition at the budding site could be connected to the ESCRT-III dependent formation of an intraluminal vesicle or to its abscission. A crucial role of lipids in MVB formation has been pointed out before (Matsuo *et al*. 2004).

## Results and discussion

### Parallel function of the Vps55/Vps68 complex and the SERINC homologue Tms1

To identify proteins that are functionally related to Vps55/Vps68 we consulted the “Yeast Fitness Database” (Hillenmeyer *et al*. 2010). There the growth fitness of deletion strains from the yeast whole-genome deletion collection in the presence of various chemicals was quantified.

For each pair of deletions, the fitness profiles across all experiments were compared and a co-fitness score was calculated, which varies between -1 (no correlation at all, opposite phenotype) and +1 (perfect correlation, identical phenotype). The seven top scoring interactors for *VPS68* (correlation >0.5) are listed in Fig. 1A (description of the genes in Tab.S2). The validity of the dataset is underscored by the finding that the *VPS55* deletion was the top scoring interaction (correlation=0.644). Second on the list of top scoring interactors is the ESCRT-III protein Did2, which could be involved in the Vps55/Vps68/ESCRT-III interaction. Another ESCRT-protein on the list is Mvb12, a member of ESCRT-I. We also examined the top scoring deletions for co-fitness among each other. Intriguingly, the deletions form a highly interconnected network of co-fitness interactions (Fig. 1B).

**Figure 1:**
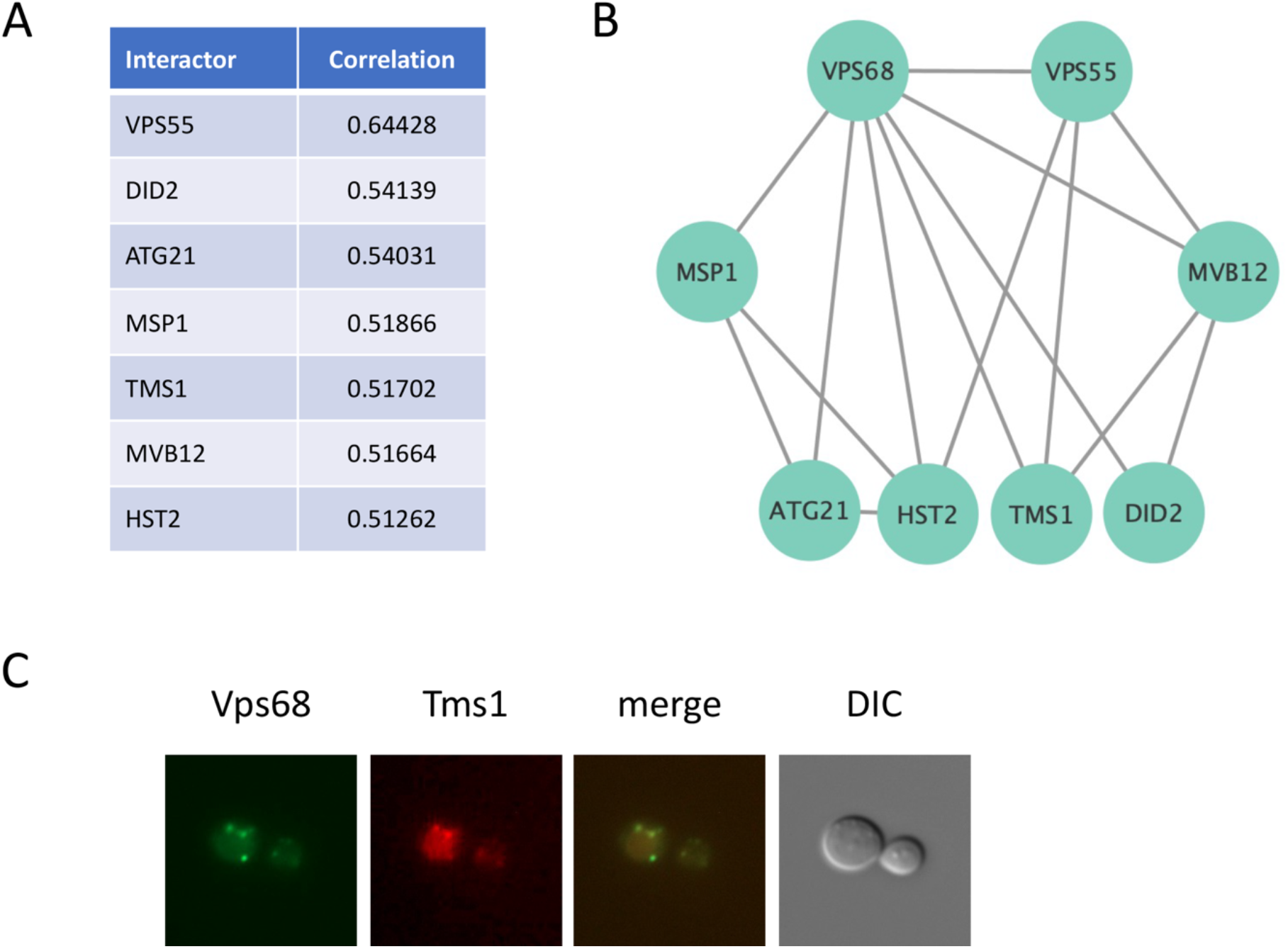
Co-fitness interactions between the *VPS68* deletion and other deletion mutants. (A) The top scoring interactors with the *VPS68* deletion obtained from the Yeast Fitness Database (Hillenmeyer *et al*. 2010). Listed are the deletion mutants with a co-fitness score above 0.5. (B) Co-fitness interactions with a co-fitness score of at least 0.4 among the top scoring interactors of *Δvps68*. (C) Tms1 co-localizes with Vps68. The localization of sfGFP-Vps68 and Tms1-mCherry in strain RKY3653 was examined by fluorescence microscopy.

The SERINC homologue Tms1 was selected for further study, since it bears some resemblance with the Vps55/Vps68 complex (Fig. S1). The core structure of the SERINC protein family is formed by two blocks of four transmembrane domains (TMDs) (Pye *et al*. 2020; Leonhardt *et al*. 2023) and Vps55 and Vps68 consist of four putative TMDs each. So far, no phenotype could be detected for the *TMS1* deletion (Grossman *et al*. 2000). First the intracellular localization of Tms1 was examined by fluorescence microscopy with gene fusions integrated into the yeast genome. With Tms1-mCherry about half a dozen dots were observed, which perfectly colocalized with sfGFP-Vps68 dots (Fig. 1C). This indicates that Vps55/Vps68 and Tms1 function at the same site in the cell. The dots most likely correspond to endosomal structures.

Next, the effect of the *VPS68* and *TMS1* deletions on the intracellular localization of the endocytic cargo protein Ste6 was examined (Fig. 2). In the wildtype strain Ste6-GFP staining was mostly detected in the lumen of the vacuole, reflecting the high constitutive turnover of Ste6 (KÖLLING AND HOLLENBERG 1994). Occasionally, a dot-like structure was observed next to the vacuole (0.04 dots/cell). In the *Δvps68* and *Δtms1* single mutants basically the same pattern was seen, but with a higher number of dots (0.24 and 0.26 dots/cell). In the double mutant the number of dots was increased to 1.4 dots/cell. This localization pattern is different from the ones previously described for other vacuolar protein sorting (*vps*) mutants (Raymond *et al*. 1992). According to this classification, Ste6-GFP neither displays a class D nor a class E localization. In class D mutants Ste6 shows enhanced recycling to the bud cell surface (Krsmanovic′ *et al*. 2005) and in class E mutants it accumulates in a brightly staining dot next to the vacuole and in the limiting membrane of the vacuole (Kranz *et al*. 2001). In the *Δvps68 Δtms1* mutant just the number of late endosomes appears to be increased.

**Figure 2:**
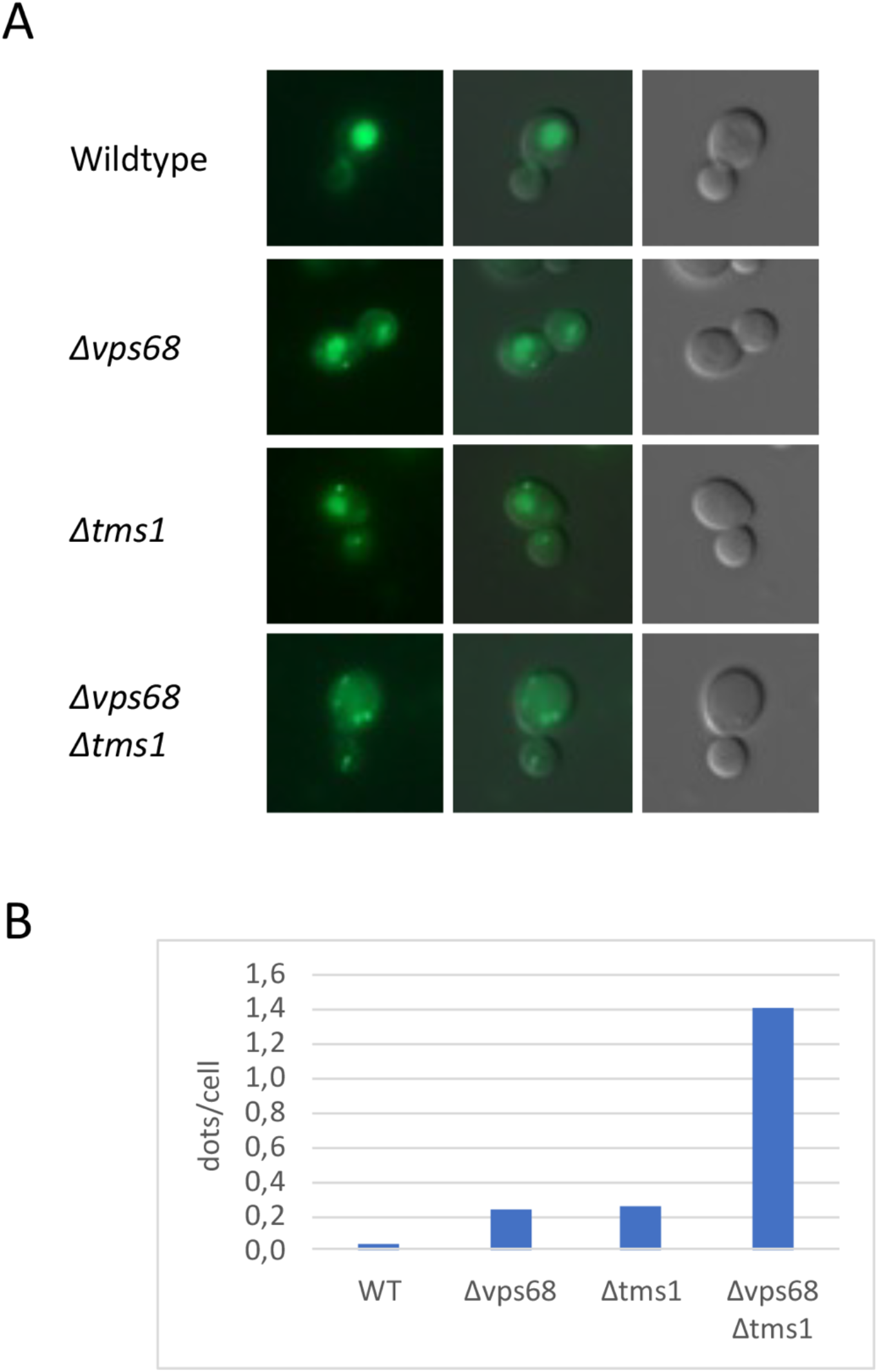
Effect of *TMS1* and *VPS68* deletions on the intracellular localization of Ste6. (A) The localization of Ste6-GFP expressed from the single copy plasmid pRK744 was examined in the different mutant strains by fluorescence microscopy. From top to bottom: RKY3319 (WT), RKY3320 (*Δvps68*}, RKY3659 (*Δtms1*), RKY3660 (*Δtms1 Δvps68*). (B) Quantification of the number of endosomal dots per cell (at least 100 cells were counted in each case).

We were then interested to see, whether this altered localization had an impact on the turnover of the Ste6 protein. The short-lived endocytic cargo protein Ste6 is transported to the yeast vacuole for degradation via the multivesicular body pathway (KÖLLING AND HOLLENBERG 1994). As described previously, Ste6 is moderately stabilized in the *Δvps68* mutant (ALSLEBEN AND KÖLLING 2022). To see whether Tms1 is involved in endocytic trafficking similar to Vps68, we examined its role in Ste6 degradation by a gal-depletion experiment (Fig. 3). Cells containing *STE6* under the control of the *GAL1* promoter were grown on galactose containing medium to induce expression of *STE6* and were then shifted to glucose containing medium to repress expression from the *GAL1* promoter. Then equal aliquots of cells were collected at time intervals and examined for the amount of Ste6 present (Fig. 3). The *TMS1* deletion displayed the same degree of stabilization as the *VPS68* deletion (half-lives 26 min and 27 min, respectively, compared to 14 min for the wildtype). Thus, both deletions moderately affect the half-life of Ste6 in a similar manner. However, when both deletions were combined, the Ste6 protein was completely stable (half-life 173 min). This pronounced synthetic phenotype indicates that the Vps55/Vps68 complex and Tms1 perform the same, parallel function at endosomes.

**Figure 3:**
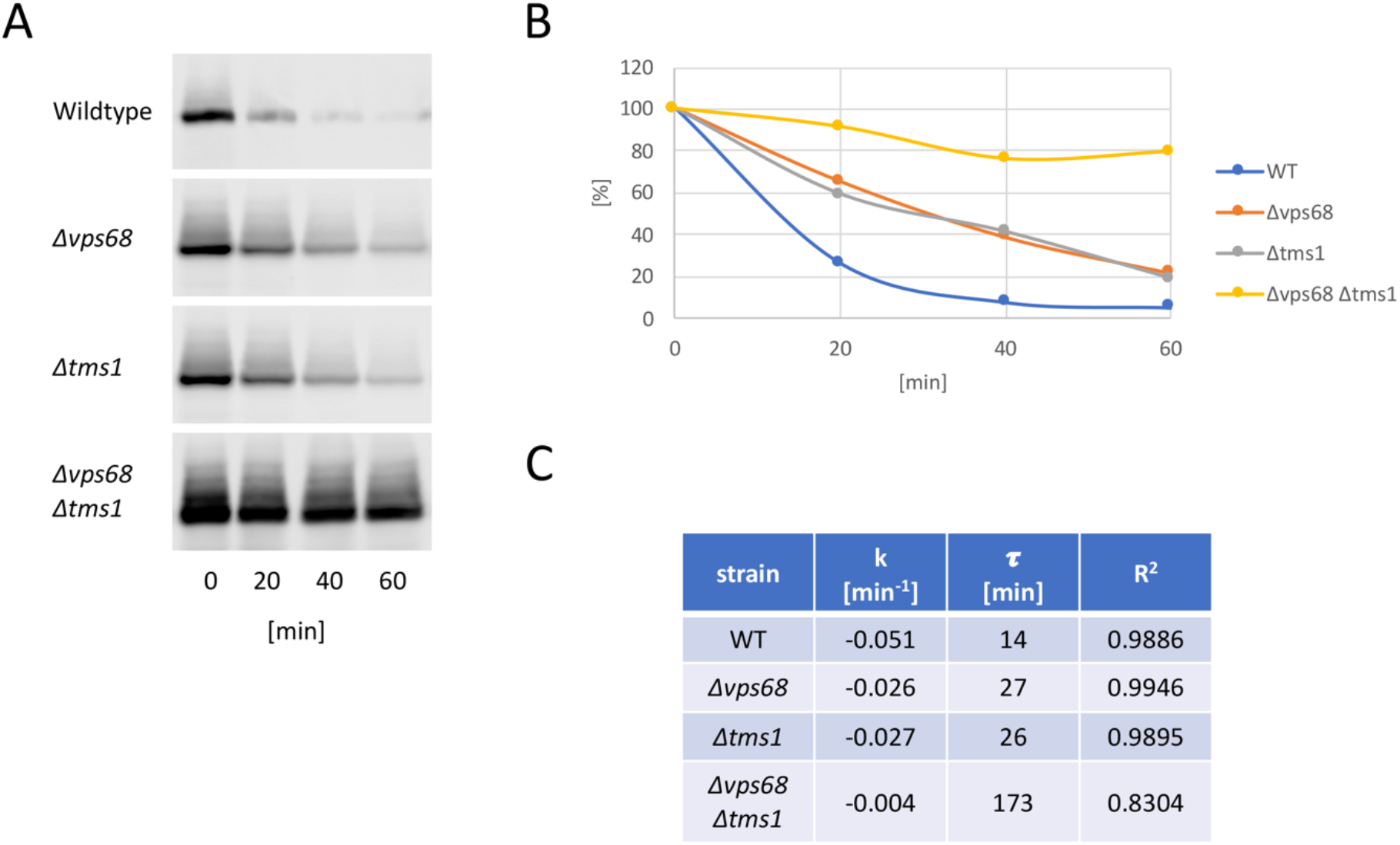
Ste6 is strongly stabilized in a *Δtms1 Δvps68* double mutant. The Ste6 turnover was investigated by a gal depletion experiment. (A) Equal aliquots of cell culture were collected at time intervals and examined for Ste6 by western blotting. The collection of samples was started 20 min after *GAL1* promoter shut-off. From top to bottom: RKY3319 (WT), RKY3320 (*Δvps68*}, RKY3659 (*Δtms1*}, RKY3660 (*Δtms1 Δvps68*). (B) The western blot signals were quantified by ImageJ. (C) The rate constant (k), the half-life (τ) and the correlation coefficient (R^2^) were calculated for the strains from the data in (B). Shown is one out of two independent experiments, which gave similar results.

### Vps55 and Tms1 interact with ESCRT-III

In our previous study we identified Vps68 as an interaction partner of ESCRT-III (ALSLEBEN AND KÖLLING 2022). Since Vps68 forms a complex with Vps55 (Schluter *et al*. 2008), we were interested to see, if Vps55 as well associates with ESCRT-III. For detection Vps55 was tagged with superfolder GFP at its N-terminus (sfGFP-Vps55). Vps55 was immunoprecipitated from cell extracts with anti-sfGFP antibodies and the immunoprecipitates were analyzed for the presence of ESCRT-III proteins by western blotting (Fig. 4A). All tested ESCRT-III proteins could be specifically co-immunoprecipitated with sfGFP-Vps55. The co-IP pattern was in line with our previous observations (Heinzle *et al*. 2019). Did2 turned out to be the most abundant ESCRT-III protein in the precipitated complexes, while the core ESCRT-III components Snf7, Vps2 and Vps24 were present in about equal amounts with a slight preference for Vps2 (Fig. 4C). In addition, a small amount of Mos10 was detected in the complexes.

**Figure 4:**
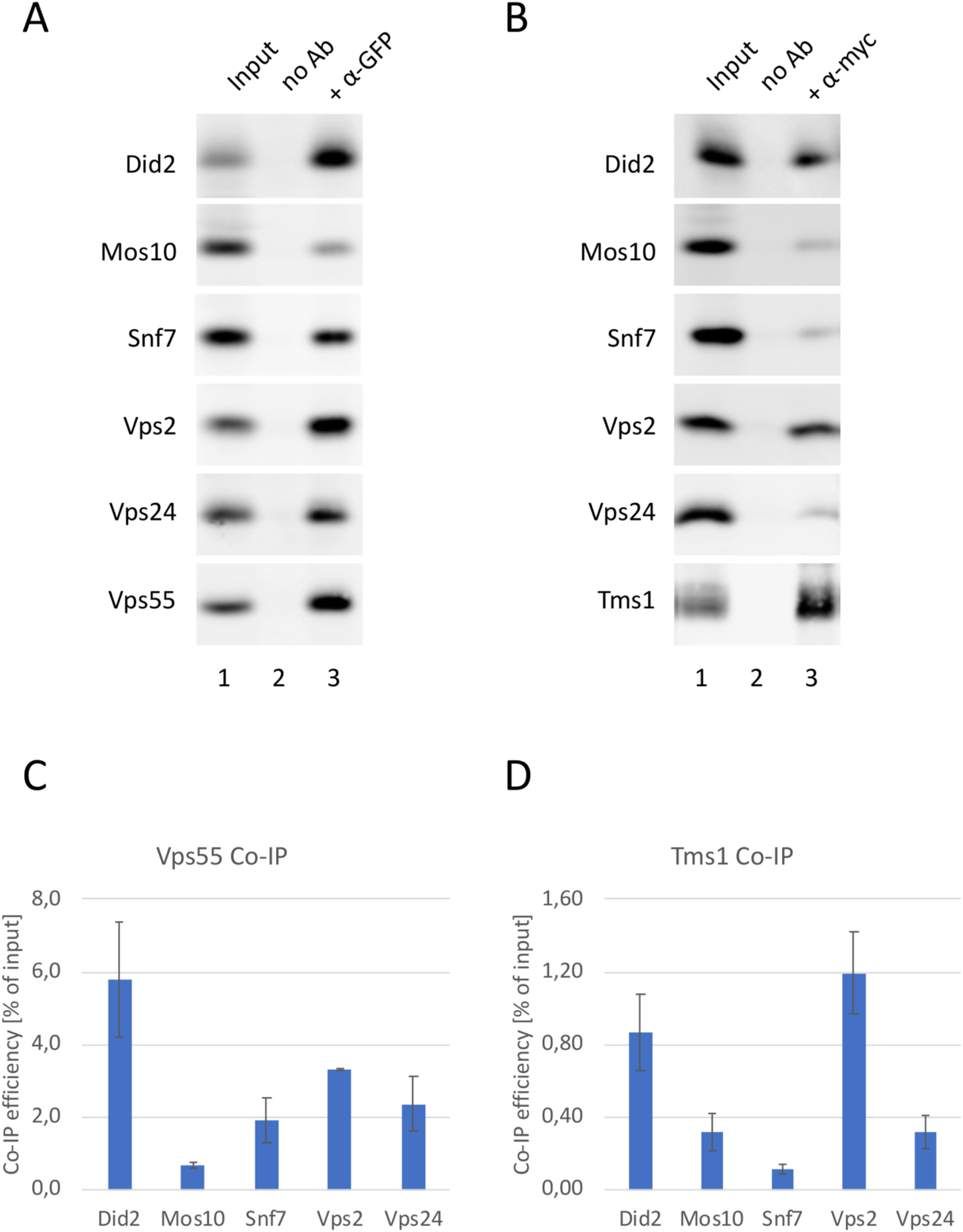
Vps55 and Tms1 interact with ESCRT-III. Immunoprecipitation of sfGFP-Vps55 from strain RKY3427 with anti-GFP antibodies (n=2) (A) and immunoprecipitation of Tms1-13myc from RKY3726 with anti-myc antibodies (n=3) (B). The immunoprecipitates were analyzed for the presence of ESCRT-III proteins by western blotting with specific antibodies against the different ESCRT-III proteins as indicated. Lane 1: input, lane 2: IP without antibodies, lane 3: IP with antibodies (primary IPs, bottom panels: 20 % of input, co-IPs, top panels: 2 % of input). (C, D) The western blot signals were quantified with ImageJ and the co-IP efficiencies with standard deviations were calculated (% of input).

Next, Tms1 was examined for its association with ESCRT-III by co-immunoprecipitation. All tested ESCRT-III proteins could also be immunoprecipitated by Tms1-13myc (Fig. 4B). However, the Tms1 co-IP pattern (Fig. 4D) was different from the Vps55 pattern (Fig. 4C). Tms1 mainly precipitated a roughly 1:1 complex of Did2 and Vps2, while Snf7, Mos10 and Vps24 were only present in low amounts. This finding is intriguing, since we postulated before from genetic experiments that Did2 and Vps2 form a distinct complex (Brune *et al*. 2019).

The co-immunoprecipitation result suggests that Vps55/Vps68 and Tms1 are not completely interchangeable in their function. We have suggested before that ESCRT-III complexes undergo remodeling during the process of ILV formation (ALSLEBEN AND KÖLLING 2022). Especially the relationship between the Snf7 complex and the Mos10 complex is unclear at present (ALSLEBEN AND KÖLLING 2022; Pfitzner *et al*. 2023). It is possible that the complex precipitated by Tms1 represents a late stage in the ESCRT-III cycle.

In conclusion, our data establish a link between lipid remodeling and ESCRT-III function. The Vps55/Vps68 complex appears to perform the same or a similar function as the SERINC homologue Tms1. Since human SERINCs are lipid-flipping scramblases, it is likely that Vps55/Vps68 has a similar activity. Loss of this activity leads to a delay in the degradation of the endocytic cargo protein Ste6. Since Vps55/Vps68 and Tms1 directly associate with ESCRT-III, it is plausible to assume that lipid remodeling at the site of ILV formation is crucial for ESCRT-III function.

## Materials and methods

### Yeast strains and media

For immunoprecipitation experiments yeast cells were grown overnight to exponential phase in YPD medium (1 % yeast extract, 2 % peptone, 2 % glucose), for gal depletion experiments they were first grown overnight to exponential phase in YPGal medium (1 % yeast extract, 2 % peptone, 2 % galactose, 0.2 % glucose) and were then shifted to YPD medium. For fluorescence microscopy cells were grown in SD/CAS medium (0.67 % yeast nitrogen base, 1 % casamino acids, 2 % glucose, 50 mg/l uracil and tryptophan) to avoid autofluorescence from the YPD medium. For plasmid selection uracil was omitted from the medium. The yeast strains used are listed in Tab. S1. Most yeast strains are derived from JD52 (J. Dohmen, Cologne, Germany) by the integration of PCR-cassettes into the yeast genome (Longtine *et al*. 1998). Strain RKY3726 was derived from YWO 0340 (K. Kuchler, Vienna). Plasmid pRK744 is a CEN plasmid with URA3 marker carrying a *STE6-GFP* fusion under the control of the *STE6* promoter.

### Co-immunoprecipitation

For co-immunoprecipitation (co-IP) experiments, 50 OD_600_ cells of an exponential YPD culture were harvested, washed with PBS and resuspended in 200 μl PBS with protease inhibitors. The cells were lysed by glass-beading for 5 min at 4°C. After lysis, 600 μl PBS were added and the solution was spun at 500 g for 5 min to remove cell debris. Then 2.5 mM of the cleavable crosslinker DSP (dithiobis (succinimidylpropionate)) (Thermo Scientific, Waltham, USA) were added to the supernatant and the solution was incubated for 30 min at RT on a rocker. The crosslinking reaction was stopped by quenching with 100 mM Tris-HCl pH 8.0 and the membranes were dissolved by 1 % Triton X-100 for 15 min at RT on a rocker. Another 500 g spin was applied to remove any formed precipitates. Then the supernatant was incubated with anti-GFP antibodies (Nordic-MUbio, Susteren, Netherlands) or anti-myc antibodies (Dianova, Hamburg, Germany) for 1 h at 4°C on a rocker followed by an incubation with 200 μl of a 20 % slurry of protein A sepharose beads (GE Healthcare, Uppsala, Sweden) for 1 h at 4°C. The solution was washed three times with 1 ml PBS at 500 g for 1 min. The beads were resuspended in a mixture of 100 μl PBS and 100 μl SDS sample buffer and heated to 95°C for 5 min. Then the immunoprecipitates were analyzed for the presence of ESCRT-III proteins by western blotting with specific antibodies.

### Fluorescence microscopy

Yeast cells were grown overnight to exponential phase in SD/CAS medium. The yeast cell suspension was applied to concanavalin A coated slides and imaged with a Zeiss Axio-Imager M1 fluorescence microscope equipped with an AxioCam MRm camera (Zeiss, Göttingen, Germany). Images were acquired with the Axiovision Software and processed with Photoshop Elements.

## Data availability

All data are available in the main text or supplementary materials.

## Acknowlegdements

I like to thank Thomas Brune for his assistance with the experiments.

## Supplemental material

**Tab. S1:**
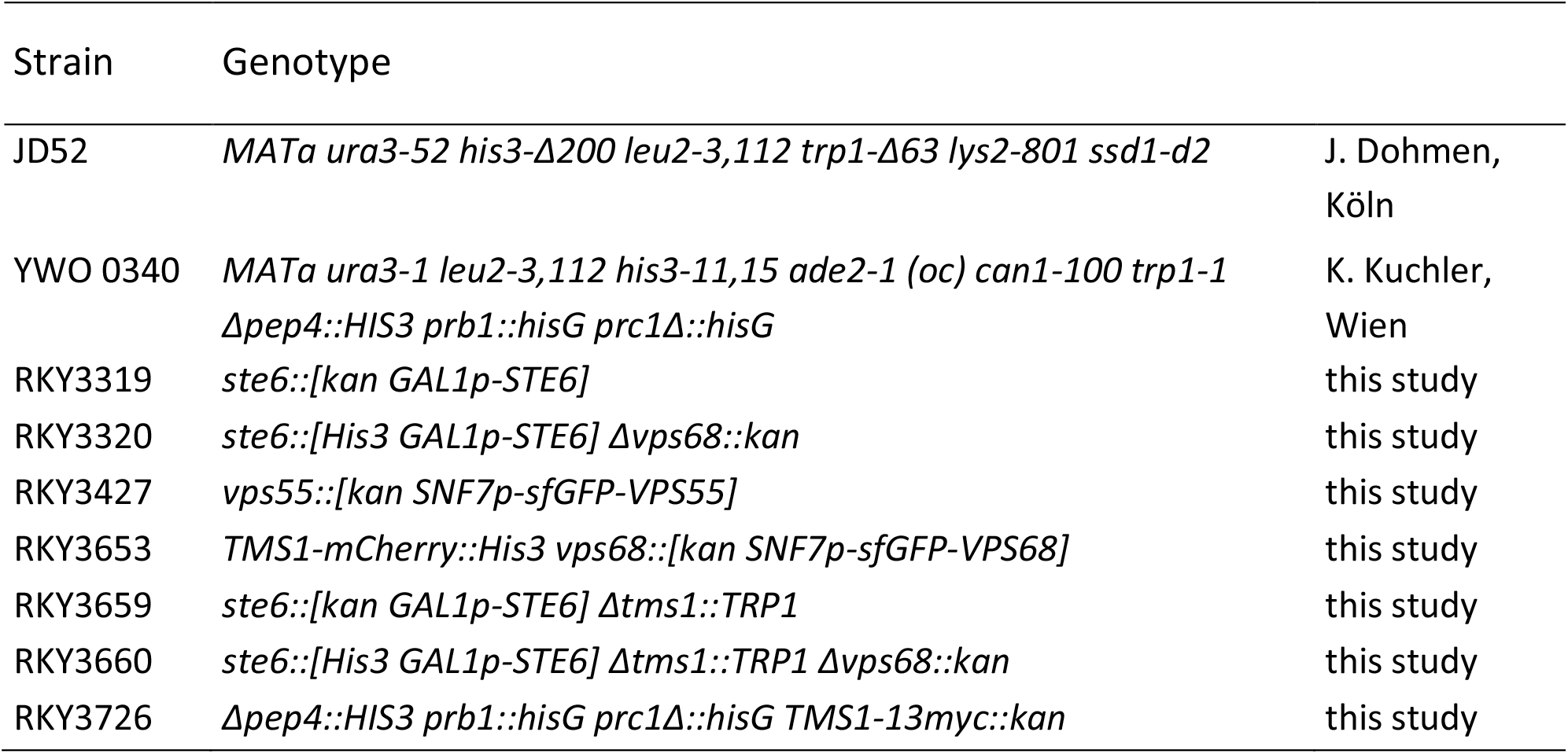
Yeast strains.

| Strain | Genotype |  |
| --- | --- | --- |
| JD52 | *MATa ura3-52 his3-Δ200 leu2-3,112 trp1-Δ63 lys2-801 ssd1-d2* | J. Dohmen,<br>Köln |
| YWO 0340 | *MATa ura3-1 leu2-3,112 his3-11,15 ade2-1 (oc) can1-100 trp1-1*<br>*Δpep4::HIS3 prb1::hisG prc1Δ::hisG* | K. Kuchler,<br>Wien |
| RKY3319 | *ste6::[kan GAL1p-STE6]* | this study |
| RKY3320 | *ste6::[His3 GAL1p-STE6] Δvps68::kan* | this study |
| RKY3427 | *vps55::[kan SNF7p-sfGFP-VPS55]* | this study |
| RKY3653 | *TMS1-mCherry::His3 vps68::[kan SNF7p-sfGFP-VPS68]* | this study |
| RKY3659 | *ste6::[kan GAL1p-STE6] Δtms1::TRP1* | this study |
| RKY3660 | *ste6::[His3 GAL1p-STE6] Δtms1::TRP1 Δvps68::kan* | this study |
| RKY3726 | *Δpep4::HIS3 prb1::hisG prc1Δ::hisG TMS1-13myc::kan* | this study |

**Tab. S2:** *Saccharomyces* Genome Database (SGD) descriptions of interactors.

| Gene | Description |
| --- | --- |
| ATG21 | Phosphoinositide binding protein; required for vesicle formation in cytoplasm-to-vacuole targeting (CVT) pathway, autophagy, micronucleophagy and mitophagy; binds to several phosphoinositides including PtdIns(3,5)P<sub>2</sub>, PI3P and PI4P; involved in PI3P-dependent recruitment and organization of both Atg12-Atg5-Atg16 complex and Atg8p at pre-autophagosomal structure; necessary for oxidant-induced cell death; targeted to vacuole via AP-3 pathway; WD-40 repeat containing PROPPIN family member |
| DID2 | Class E protein of the vacuolar protein-sorting (Vps) pathway; binds Vps4p and directs it to dissociate ESCRT-III complexes; forms a functional and physical complex with Ist1p; human ortholog may be altered in breast tumors |
| HST2 | Cytoplasmic NAD(+)-dependent protein deacetylase; deacetylation targets are primarily cytoplasmic proteins; member of the silencing information regulator 2 (Sir2) family of NAD(+)-dependent protein deacetylases; modulates nucleolar (rDNA) and telomeric silencing; possesses NAD(+)-dependent histone deacetylase activity in vitro; contains a nuclear export signal (NES); function regulated by its nuclear export |
| MSP1 | Highly-conserved N-terminally anchored AAA-ATPase; distributed in the mitochondrial outer membrane and peroxisomes; involved in mitochondrial protein sorting; functions as an extraction engine in local organelle surveillance to remove and initiate degradation of mistargeted proteins, ensuring fidelity of organelle-specific localization of tail-anchored proteins; contains an N-terminal transmembrane domain and C-terminal cytoplasmic ATPase domain |
| MVB12 | ESCRT-I subunit required to stabilize ESCRT-I core complex oligomers; the ESCRT-I core complex (Stp22p, Vps28p, Srn2p) is involved in ubiquitin-dependent sorting of proteins into the endosome; deletion mutant is sensitive to rapamycin and nystatin |
| TMS1 | Vacuolar membrane protein of unknown function; is conserved in mammals; predicted to contain eleven transmembrane helices; interacts with Pdr5p, a protein involved in multidrug resistance |
| VPS55 | Late endosomal protein involved in late endosome to vacuole transport; functional homolog of human obesity receptor gene-related protein (OB-RGRP) |

**Figure S1:**
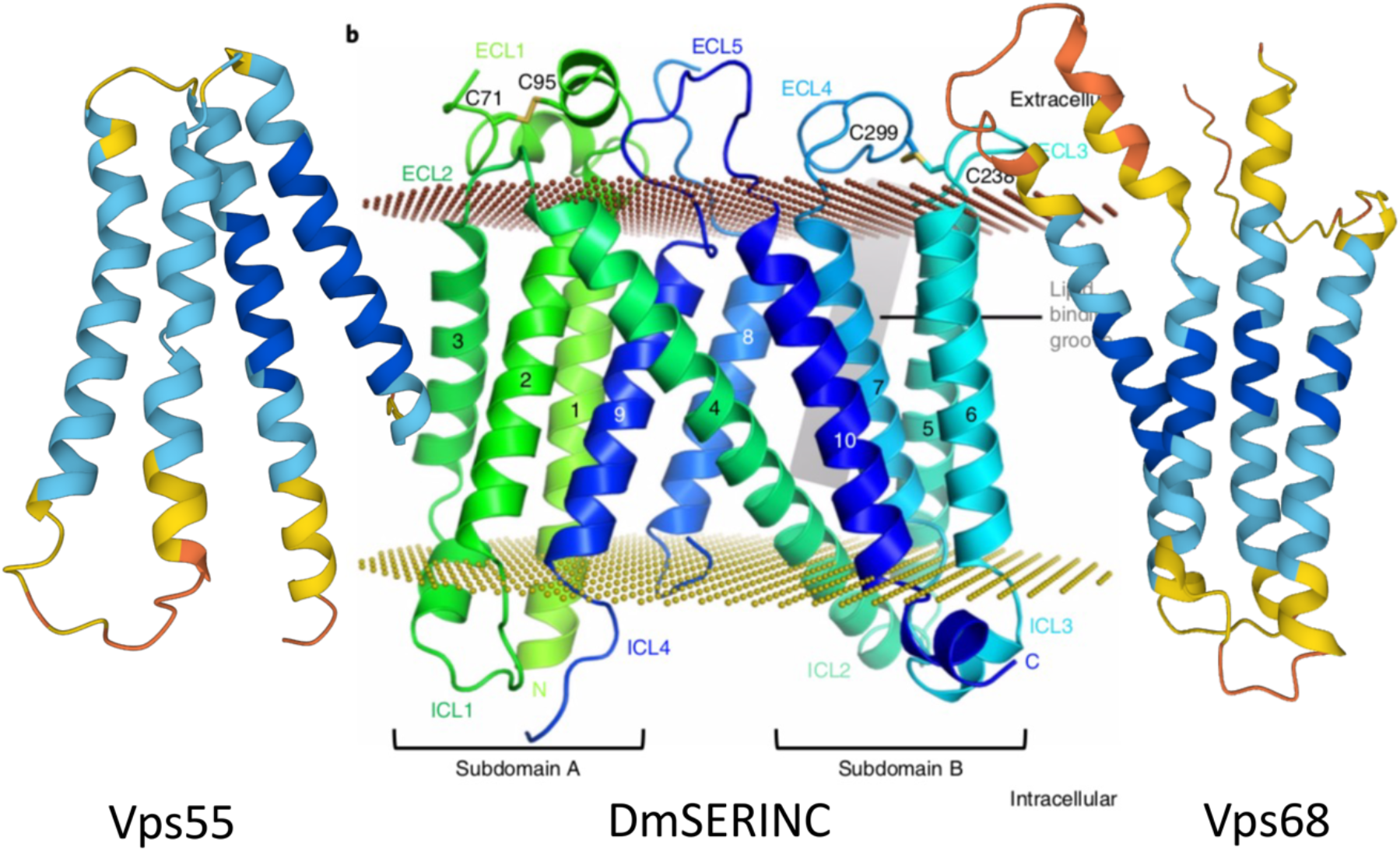
The cryo-EM structure of DmSERINC taken from (Pye *et al*. 2020) is compared with the structures of Vps55 and Vps68 as predicted by AlphaFold (Varadi *et al*. 2022).

